# scGrapHiC : Deep learning-based graph deconvolution for Hi-C using single cell gene expression

**DOI:** 10.1101/2024.02.07.579342

**Authors:** Ghulam Murtaza, Byron Butaney, Justin Wagner, Ritambhara Singh

## Abstract

Single-cell Hi-C (scHi-C) protocol helps identify cell-type-specific chromatin interactions and sheds light on cell differentiation and disease progression. Despite providing crucial insights, scHi-C data is often underutilized due the high cost and the complexity of the experimental protocol. We present a deep learning framework, scGrapHiC, that predicts pseudo-bulk scHi-C contact maps using pseudo-bulk scRNA-seq data. Specifically, scGrapHiC performs graph deconvolution to extract genome-wide single-cell interactions from a bulk Hi-C contact map using scRNA-seq as a guiding signal. Our evaluations show that scGrapHiC, trained on 7 cell-type co-assay datasets, outperforms typical sequence encoder approaches. For example, scGrapHiC achieves a substantial improvement of 23.2% in recovering cell-type-specific Topologically Associating Domains over the baselines. It also generalizes to unseen embryo and brain tissue samples. scGrapHiC is a novel method to generate cell-type-specific scHi-C contact maps using widely available genomic signals that enables the study of cell-type-specific chromatin interactions.

**Availability:** https://github.com/rsinghlab/scGrapHiC

## 1. Background

The 3D organization of the genome plays a critical role in modulating a wide range of cellular functions, including gene expression, that drive cell differentiation and disease progression [Fa95; Kl01]. Genome-wide conformation capture techniques, like Hi-C [Ra14], Micro-C [SCH22], and ChIA-PET [Li14], measure the genomic spatial interactions and offer insights into how they regulate the gene expression. These experiments produce an array of paired-end reads, where each paired-end captures two DNA sequences (or genomic loci) that interact in the 3D space. These paired-end reads are typically coalesced into a contact map of size *N* × *N*, where each entry represents a genomic region of size 1Kbp to 1Mbp. Analysis of these contact maps sheds light on important genome organizational components that are tied to gene regulation, such as A/B compartments, Topologically Associating Domains (TADs), and chromatin loops [Ra14]. Building upon these techniques, a recent extension of the Hi-C protocol is single-cell Hi-C (scHi-C) [St17]. This innovative approach provides a detailed spatial view of the genome at a single-cell resolution, allowing researchers to decipher regulatory mechanisms in each cell.

Constructing high quality Hi-C contact maps (≤ 5Kbp) requires billions of reads. Obtaining these contact maps is costly and can be infeasible when studying rarer cell types, such as some type of cancers [Dí18], where obtaining large number of cells may be impractical. Compared to bulk protocols (Hi-C, Micro-C, and ChIA-PET), scHi-C presents additional experimental challenges [GG21] resulting in sparser contact maps because the reads are further divided across various cell populations. Even though scHi-C provides exciting insights into structural rearrangements at individual cell resolution, the protocol is not as widely used due to its limitations [GG21].

To address the experimental complexity of the Hi-C protocol, several deep learning methods have been developed that generate genome-wide contact maps for bulk Hi-C by relying on easier-to-obtain genomic measurements. We call them sequence encoder methods as they encode the DNA sequence or a sequential 1D genomic measurement to predict the Hi-C contact map. Akita [FKP20] was the first sequence encoder method that predicted Hi-C contact maps using DNA sequences. However, since it did not rely on any cell-type-specific signal, Akita could not predict cell-type-specific interactions. C.Origami [Ta22], Epiphany [Ya23], and Chromafold [Ga23] have extended the sequence encoding framework of Akita to input different cell-type-specific signals (ChIP-Seq or ATAC-Seq) and predict cell-type-specific Hi-C contact maps. These sequence encoder methods make accurate predictions on bulk Hi-C datasets that capture the average cell population signal. However, their applicability is not extended to scHi-C, constraining their ability to predict the biological variations in the chromatin structure at a higher cell-level resolution, which reveals heterogeneity in the measured cell or tissue-types.

Deconvolution methods can potentially recover the cell-type-specific heterogeneity from a bulk Hi-C contact map. THUNDER [Ro22] and DECOOC [Wa23] extract cell-type specific interactions and cell population percentages from a bulk Hi-C sample. THUNDER has two non-negative matrix factorization steps. In the first step it performs feature selection on the bulk Hi-C matrix and then deconvolves cell proportions based on those selected features. DECOOC uses a convolutional neural network architecture to predict cell proportions in the bulk sample. The reliance of these methods on experimental tissue-specific bulk Hi-C contact maps severely constrains their applicability because obtaining bulk Hi-C for every tissue-type is costly and might be infeasible for certain tissues. Both methods require prior information on the number of type of cells in the bulk sample, which may not be easily available. The combined experimental and computational limitations makes it challenging to acquire and study genome-wide scHi-C contact maps and restrict a refined understanding of cell development or diseases.

We present a deep learning framework called **scGrapHiC** (Fig. 1), which is a graph deconvolution method that predicts cell-type-specific pseudo-bulk single-cell Hi-C contact maps. It uses pseudo-bulk single-cell RNA-seq as a guiding signal to extract cell-type-specific contact maps from a bulk Hi-C sample. scGrapHiC innovates over previous work by using scRNA-seq as input, which is much more widely available than other single-cell measurements. However, scRNA-seq presents unique challenges, given its weaker correlation with the 3D genomic structure than typically used measurements, like ATAC-seq or ChIP-seq, for bulk Hi-C predictions. We resolve these challenges by providing additional support through CTCF binding affinities and CpG scores and relying on the structure that we deconvolve from a bulk Hi-C contact map. Unlike existing deconvolution approaches, scGrapHiC predicts genome-wide contact maps and does not require any prior information about the number of cell-types in the bulk Hi-C sample.

**Abb. 1:**
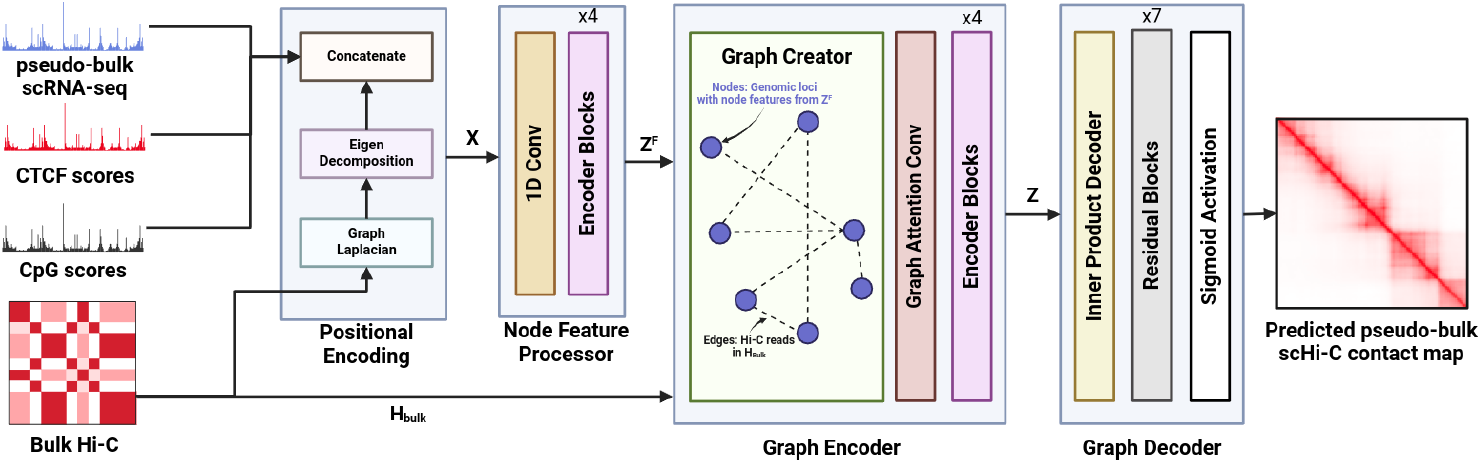
Overview of scGrapHiC. scGrapHiC is a deep learning framework that extracts cell-type specific scHi-C from a bulk Hi-C contact map using scRNA-seq as a guide signal. scGrapHiC has four major components: The first component - Positional Encoding - extracts the positional encodings from the bulk Hi-C and concatenates them with our input node feature set that contains scRNA-seq, CTCF, and CpG scores. The second component - Node Feature Processor - maps this feature set into a joint representation space. The third component - Graph Encoder - produces a node latent set that represents the likelihood of contacts being extracted via deconvolution on the bulk Hi-C using the provided joint node feature set as a guide signal. The last component - Graph Decoder - maps these likelihoods to the scHi-C output space.

We train scGrapHiC on a recently published scRNAseq and scHi-C co-assay dataset [Li23] across 7 different cell-types. scGrapHiC outperforms sequence-based baselines (for example, an improvement of 22.3% in recovering cell-type specific TADs) demonstrating the importance of the graph deconvolution from bulk Hi-C data. We show that a widely available bulk Hi-C sample (for example, from Embryonic Stem Cells (ESCs)) can be used for the deconvolution step across cell-types and tissue samples. Finally, we show that scGrapHiC generalizes to unseen embryo and brain tissue scRNA-seq inputs without any retraining or finetuning, highlighting that it can be adapted to a wide range of use cases.

Overall, scGrapHiC is a novel approach to predict pseudo-bulk scHi-C that also generalizes to unseen cell types using a widely available pseduo-bulk scRNA-seq. Leveraging easily accessible input datasets, scGrapHiC can enable the study of the 3D genomic organization at a finer cellular resolution for the research community.

## 2 Methods and Materials

As shown in Fig. 1, scGrapHiC predicts pseudo-bulk scHi-C contact maps from pseudo-bulk scRNA-seq by relying on deconvolved structure from bulk Hi-C, CTCF, and CpG to provide additional structural support. Each input corresponds to a genomic region binned at 50Kbp resolution to predict the scHi-C contact map corresponding to that region at the same resolution. For the inputs, each bin or the genomic locus either contains the combined observed gene expression values for pseudo-bulk scRNA-seq or the average scores for CTCF and CpG modalities. scGrapHiC achieves this through four main components:

### 2.1 Positional Encodings module

Existing sequence encoding methods assume a sequential structure on the input genomic measurement, even though the measurements originate from complex and 3D structured chromatin. Such sequential modeling ignores the relative positions of genomic loci in 3D space. Importantly, this positioning encodes a prior distribution on how likely two genomic loci might interact structurally, for instance regions of DNA in the same A/B compartments or TADs are more likely to interact and be functionally related [Ra14]. We learn this distribution by extracting graph positional encodings through bulk Hi-C data. We compute these encodings by first constructing a normalized Laplacian matrix *L*_norm_ from an input bulk Hi-C contact map *H*_*bulk*_ as follows:

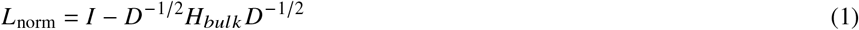

Here, *D* is the degree matrix of *H*_*bulk*_ that contains the the number of edges attached to each node and *I* is the identity matrix [Dw20]. We decompose the Laplacian matrix *L*_norm_ into its eigenvectors. These eigenvectors, extracted from the bulk Hi-C, capture the underlying hierarchical topology of the DNA. Finally, we concatenate the top *k* (*k* = 16) components of the eigenvectors with our node feature vector *X*, comprised of CTCF and CpG scores and pseudo-bulk scRNA-seq reads. Our calculated encodings allow us to enrich *X* with the prior that genomic loci belonging to the same A/B compartment or TADs are more likely to interact [Di12; Li09]. Conversely, genomic loci that are farther apart in the 3D space are less likely to interact [Ra14]. Thus, nodes representing these loci have similar or dissimilar values, respectively.

### 2.2 Node Feature Processor module

Genomic features tend to interact with each other at both local and global scales. Therefore, instead of treating these features independently, we explicitly incorporate the local and global interactions in scGrapHiC through the Node Feature Processor module. First, we apply a single Conv1D filter with a window size of 16 on the input node feature set *X*, which extracts a localized feature set *F*. Specifically, *F* encodes the relationships between CTCF and CpG scores, structural neighborhood, and the gene expression profile within the ‘local’ Conv1D window. The Conv1D allows scGrapHiC to identify strong genomic signals over potential background noise, thus acting as a learnable post-processing step. Next, we pass the feature set *F* through a series of Transformer encoder blocks, which capture the long-range or ‘global’ interactions between all features learned in the previous step. At the core of the transformer encoder block is the self-attention operation [De18]. The self-attention operation produces an updated node feature set *Z* which, for each loci *i*, encodes the contributions of all other node features in *F* scaled by their relative relevance to features in node *i*. This captures the global interactions of the feature in node *i* with all other nodes. The self-attention score is calculated as follows:

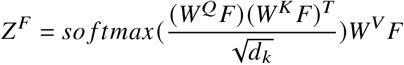

Here *W*^*Q*^, *W*^*K*^, *W*^*V*^ are learnable parameters and *d*_*k*_ is a parameter that scales the output of the dot products for stable training. The output vector *Z* ^*F*^ from the Node Feature Processor module is an enriched node feature set *X* that captures the local and global interactions among genomic features.

### 2.3 Graph Encoder module

The approach of relying on genomic measurements, such as ATAC-seq to predict bulk Hi-C is impractical for predicting scHi-C due to the technical constraints of the single-cell protocol. Existing single-cell experiments can perform either a chromosome capture assay (Hi-C) or an ATAC-seq experiment at a given time because they both rely on accessing DNA, making it impractical to acquire input-output pairs to train a deep-learning based model. The availability of scRNA-seq measurements makes it a good choice for scGrapHiC to capture cell-type specificity of scHi-C data. However, because it is an indirect measurement of the genome, the scRNA-seq signal tends to be weakly correlated with the structure of the genome. Therefore, we deconvolve additional structural support from bulk Hi-C to allow us to map scRNA-seq to scHi-C accurately.

The Graph Encoder performs two operations - (1) deconvolution and (2) mapping. *Deconvolution* inputs the bulk Hi-C contact map and extracts the relevant structure from it. First we construct a Hi-C graph *G* = (*V, E*) with nodes *V*, where each node *v*_*i*_ has a node feature 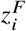 and edges *e ∈ E* from bulk Hi-C *H*^*bulk*^ connecting them. Every node *v*_*i*_ *∈ V* corresponds to the processed features of a genomic locus *i*, and an edge *e*_*i, j*_ *∈ E* is the Hi-C read observed between genomic locus *i* with another locus *j* in *H*^*bulk*^. Next, we input the graph *G* into a Graph Attention (GAT) neural network [BAY22] layer. GAT operation has two steps that work as follows:

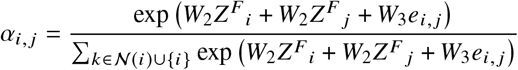

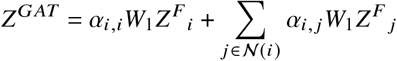

Where *W*_1_, *W*_2_ and *W*_3_ are learnable parameters. GAT aggregates the contributions of all the first-order (𝒩_*i*_) connected node features scaled by an attention score *a*_*i, j*_ . The attention score *a*_*i, j*_ represents the relevance of that target node *j* to the source node *i* based on their node features and the edge *e*_*i, j*_ connecting them. Interestingly, since *a*_*i, j*_ can prune the non-informative edges in the Hi-C graph, it extracts the relevant structure from the bulk Hi-C to predict cell-type specific scHi-C contacts using cell-type specific node features *Z* ^*F*^. These attention coefficients can effectively be seen as a deconvolution process. GAT creates the node latent *Z*^*GAT*^ by aggregating the node features of all the first order neighbors for node *i ∈* 𝒩 (*i*) scaled by the attention coefficients *a*_*i, j*_ to capture the complex interplay of cell-specific signals in node features and the deconvolved structure in the attention coefficients.

In the *mapping* phase, we map *Z*^*GAT*^ to a node latent space *Z*, which is an information-rich latent space conditioned on cell-type specific features from scRNA-seq and structural support through CTCF and CpG scores as well as deconvolved bulk Hi-C data. To acquire *Z* we apply a series of stacked transformer encoder blocks to capture all the interactions between all pairs of *i, j* in *Z*^*GAT*^ . This *Z* is learned such that an inner product of *z*_*i*_ and *z* _*j*_ would reflect the contact likelihood of genomic loci *i* and *j* in the pseudo-bulk scHi-C data.

### 2.4 Graph Decoder module

The purpose of the Graph Decoder module is to convert *Z* to the cell-type specific scHi-C contact maps. To achieve that, Graph Decoder first applies an inner product on the node latent embeddings *Z* through *⟨Z, Z*^*T*^*⟩*, which outputs a 2D contact map that contains the likelihood of spatial interaction between all pairs of genomic loci. To map these likelihoods onto the output scRNA-seq space, we then apply stacked residual blocks with a final sigmoid activation layer that produces our goal pseudo-bulk scHi-C contact map corresponding to the input scRNA-seq data in *X*.

### 2.5 Implementation details

The entire pipeline is implemented in Python (version 3.9.0), and scGrapHiC is implemented with Pytorch Lightning (version 2.1.3) using the Pytorch (version 2.1.0) and Pytorch Geomtric (version 2.3.0) backends. scGrapHiC takes in a 128 ×128 bulk Hi-C contact map with 50Kbp bin size corresponding to a 6.4 Mbp genomic region. scGrapHiC also requires a 128 × 5 node feature vector for the same region. This node feature vector contains positive and negative strands for CTCF scores and scRNA-seq reads concatenated with the CpG score vector. We combine a positional encoding vector of 16 with the node feature vector. scGrapHiC produces a 128 × 128 scHi-C output, which we compare against a target scHi-C using an MSE loss to optimize the weights of our model across 300 epochs. We show the loss curve for scGrapHiC in supplementary Fig. S1 across five different random seeds to highlight our training process is robust to initial weights. We input 6.4 Mbp genomic regions to scGrapHiC with a stride of 16, and we average predictions on all the overlapping regions to construct our final scHi-C intra-chromosomal maps. We chose to input only 6.4 Mbp genomic regions because it allowed us to maximize the biologically informative interactions we consider [Zh18] for each prediction while minimizing the sparsity in our target scHi-C contact map for stable training. We summarize the model hyperparameters and their corresponding values in the supplementary Table. S1.

## 3. Experimental Setup

### 3.1 Datasets

#### 3.1.1 Single cell datasets

We obtained single-cell datasets from a scRNA-seq and scHi-C co-assay method, HiRES [Li23]. The study (GSE223917) produced data for over 7000 mouse embryos, spanning embryo stage E7.0 to EX15, along with 400 cells from mouse brain frontal cortex. Across all the tissues and stages, we pseudo-bulked single cells using the provided labels in the metadata ^3^, and excluded all the cell types that had less than 190 cells. We chose this cutoff because it allowed us to minimize the inherent sparsity in single-cell data and maximize our coverage across all the cell types provided in the HiRES dataset. We show all the selected cell types with corresponding stage, tissue, and the observed cell count in Table 1.

**Tab. 1:** Summary of the HiRES dataset after the pseudo-bulking and filtration step. We use the embryo stages E70, E75, E85, and EX05 cell lines for training. Note that we use all chromosomes except 7 and 11 for the training set; hence, all the results capture a cross-chromosome evaluation scenario with chromosomes 7 and 11 as test set. Furthermore, we separate out EX15 and brain tissue samples to test our model’s performance in cross-chromosome, cross-tissue and embryo stage evaluations.

|  | Tissue | Stage | Number of cells |
| --- | --- | --- | --- |
| **Epiblast and PS** | Embryo | E70 | 194 |
| **Early Mesoderm** | Embryo | E75 | 204 |
| **ExE Ectoderm** | Embryo | E75 | 256 |
| **ExE Endoderm** | Embryo | E75 | 253 |
| **Neural Endoderm** | Embryo | E75 | 390 |
| **Blood** | Embryo | E80 | 233 |
| **Mix Late Mesenchyme** | Embryo | EX05 | 391 |
| **Early Neurons** | Embryo | EX15 | 255 |
| **Mix Late Mesenchyme** | Embryo | EX15 | 403 |
| **Ex1** | Brain | N/A | 203 |

We mapped the scHi-C reads of all the cells belonging to the same cell type to the mm10 genome reference at 50Kbp resolution using Cooler tools [AM20]. We discarded all inter-chromosomal reads as they tended to be extremely sparse, and we also removed X, Y, and MT chromosomes. Similarly, for scRNA-seq, we reverse mapped cell-by-gene UMI counts of all cells belonging to the same cell type back to both positive and negative strand genome track using the GENCODE mm10 (vM23) GTF file. We binned reads on both genomic tracks to a 50Kbp resolution by averaging the expression in each bin. The number of cells per pseudo-bulk varies from 190 to 400. To ensure similar coverage across all cells, we perform a library size normalization for both scRNA-seq and scHi-C:

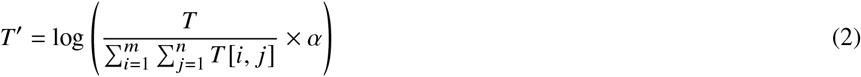

The library size normalization first computes the likelihood of observing a read at all loci in *T* . It samples a new *T* ^′^ first by multiplying by library size parameter *α* (we use a value of 25000), followed by a log scaling. Intuitively, this normalization ensures similar read distribution profiles across all pseudo-bulk cell types with different numbers of cells. We perform an additional denoising step for scHi-C because of the poor signal-to-noise ratio, which can be further amplified due to library size normalization. We first decompose the scHi-C matrix into eigenvectors *P* and their corresponding eigenvalues Λ. We soft-threshold the eigenvalues to Λ^′^ with threshold value *t* of 0.5:

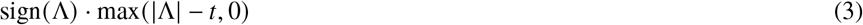

The soft-thresholding function sets all the eigenvalues smaller than the threshold value *t* to zero. Since small eigenvalues and their corresponding eigenvectors (high-frequency components) are related to experimental noise in Hi-C [Ya17], we omit their contribution by setting them to zero. We reconstruct the denoised Hi-C matrix with the soft-thresholded eigenvalues Λ^′^ by P · Λ^′^ · *P*^*−*1^. We perform an additional min-max normalization of both scRNA-seq and scHi-C to project all values between the range 0 *−* 1, which improves the stability of training process. We keep chromosomes 7 and 11 for testing purposes because they are the most gene-dense mouse chromosomes [Ma05], and we use the rest of the chromosomes for training our model. Additionally, we kept Ex1 (Exon) cells from the brain and Mixed Late Mesenchyme and Early Neuron cells from embryo stage EX15 separate for evaluations in cross-tissue and cross-embryo developmental stage evaluations.

#### 3.1.2 Bulk Hi-C datasets

Bulk Hi-C datasets were collected from a study with GEO Accession GSE82185, which contains bulk Hi-C measurements of mouse embryos’ from stage E0.5 to E4.5 as well as mouse brain cortex samples. We used the bulk Hi-C data from embryo stages E0.5 to E4.5 to identify a candidate dataset to use as an input for scGrapHiC for predicting chromatin structure of various cell-types and tissue samples. We processed the bulk Hi-C datasets similar to pseudo-bulk scHi-C and scRNA-seq data. We also include a brain cortex bulk Hi-C dataset to test whether scGrapHiC can capitalize on a tissue Hi-C measurement to accurately deconvolve scHi-C contact maps for cell types belonging to that tissue. We bin the bulk Hi-C data at a 50Kbp resolution and divide it into sub-matrices of size 128 × 128 with a stride of 32. Given the substantially deeper coverage of bulk Hi-C datasets, we do not denoise our Hi-C matrices and only perform library size normalization. Similar to scHi-C and scRNA-seq, we also min-max normalize the bulk Hi-C data. scGrapHiC uses mouse embryonic stem cells (mESC) bulk Hi-C contact maps as input unless stated otherwise.

#### 3.1.3 CTCF and CpG scores

We obtained the CTCF motif scores from CTCF R package [Do22], which receives these motif scores by scanning all three JASPAR [Ca21] CTCF PWMs in genomic DNA sequences using FIMO [GBN11]. We acquire CpG frequency using pycoMeth [Le20]. PycoMeth extracts putative CpG islands in a reference DNA sequence and generates an associated CG dinucleotide frequency for each island. Similar to the single cell and bulk data, we bin CTCF motifs and CpG scores in a 50Kbp resolution, and we normalize both CTCF and CpG scores to be in the range of 0 to 1 using a min-max scaling approach.

### 3.2 Baselines

To our knowledge, no existing methods predict genome-wide pseudo-bulk scHi-C using scRNA-seq as the input. We compare scGrapHiC against a comprehensive set of ablation models used as baselines, which capture the methodological essence of Epiphany, Chromafold and C.Origami that use only genomic sequences to predict Hi-C contact maps.

#### 1. Bulk only model

is the implementation that maps an input bulk Hi-C graph to scHi-C without any guiding cell-type-specific signal.

#### 2. scRNA-seq only model

resembles the sequence encoder methods such as Epiphany [Ya23] that relies on cell-type-specific sequential information through histone modification and CTCF ChIP-seq to predict Hi-C contacts.

#### 3. scRNA-seq+CTCF model

encompasses a set of methods like Chromafold [Ga23] and C.Origami [Ta22] that require ATAC-seq with additional structural support through CTCF motif scores to predict cell-specific bulk Hi-C contact maps.

#### 4. scRNA-seq+CTCF+CpG model

extends scRNA-seq+CTCF with additional support that provides CpG frequencies as another feature. Through this version, we investigate how additional cell-agnostic features improve performance.

#### 5. scGrapHiC

implements the graph deconvolution methods that extract the cell-typespecific scHi-C from the input bulk Hi-C graph by utilizing scRNA-seq as the guiding signal.

We exclude THUNDER and DECOOC from our baselines because the output produced by these methods is cell type proportions, which is incompatible with genome-wide contact maps scGrapHiC generates.

### 3.3 Evaluation Metrics

We evaluate the performance of predicted scHi-C contact maps with the target scHi-C contact maps using the following metrics:

#### GenomeDISCO

(GD) is a Hi-C similarity metric that models Hi-C data as a graph and compares the transition matrices at increasing timesteps *t* to compare the hierarchical organization of genome across two Hi-C contact maps [Ur18]. GD produces a score between -1 and 1, 1 representing an identical hierarchical structure between the input Hi-C matrices.

#### Stratum-Adjusted Correlation Coefficient

(SCC) is a Cochran-Mantel-Haenszel (CMH) statistic that compares the similarity of two variables while a third variable stratifies them. We use an implementation of this statistic for scHi-C matrices known as HiCRep [Ya17], which compares the similarity of Hi-C reads stratified by their distance from the matrix diagonal. HiCRep produces a correlation score between *−* 1 and 1 with a score of 1 suggesting identical contact maps.

#### Topologically Associating Domains Similarity

(TAD Sim) To compare the TAD boundaries between two Hi-C contact maps to compare the biological similarity of two Hi-C contact maps. We first call TADs in both the input and the target Hi-C contact using Chromosight [Ma20] and count the TAD boundaries that overlap as True Positives (TP). We also count False Positives (FP) features in the generated scHi-C contact map but not in the target and False Negatives (FN) as features in the target scHi-C contact map but not in the generated scHi-C contact map. We compute an F1 score as follows to quantify the TAD similarity of two scHi-C contact maps:

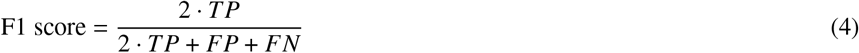

## 4. Results

### 4.1 scGrapHiC accurately predicts a pseudo-bulk scHi-C contact map using pseudobulk scRNA-seq, bulk Hi-C, CTCF, and CpG scores

An important distinction of scGrapHiC from existing sequence-based encoder methods is the use of bulk Hi-C data that it deconvolves for improved scHi-C prediction. We conduct an ablation analysis to highlight the necessity and effectiveness of this graph deconvolution approach in predicting pseudo-bulk scHi-C contact maps from scRNA-seq data. We include three sequence-based encoding approaches that capture the essence of existing encoder methods Chromafold, C.Origami and Epiphany for bulk Hi-C prediction. We also compare scGrapHiC ‘s performance against the *Bulk only model* that maps bulk Hi-C to scHi-C contact matrices. We show that *scRNA-seq only model* is insufficient to predict the single-cell structure. We further show that adding support through cell-agnostic modalities such as CTCF and CpG scores to scRNA-seq information, similar to Chromafold and C.Origami, improves the performance but it still lags behind our approach. Qualitative evaluations shown in Fig. 2 compare the imputed Hi-C contact maps for region chr7:32Mbp-38.4Mbp of Epiblast and PS cell type; we picked this cell-type because it has the smallest number of cells (194) and we chose this region because it shows a high density of chromatin regions including TADs and chromatin loops. The *Bulk only model* collapses and produces the same output for all bulk Hi-C inputs showing that we require cell-type-specific information to be able to infer any structure. The *scRNA-seq only model* can only predict the higher order structure of the chromatin, and additional support through CTCF and CpG scores improves the quality as they can predict more and sharper structural features. However, as shown in the region highlighted with a blue rectangle, none of the sequence encoder methods can predict the sub-TAD structure that scGrapHiC predicts successfully.

**Abb. 2:**
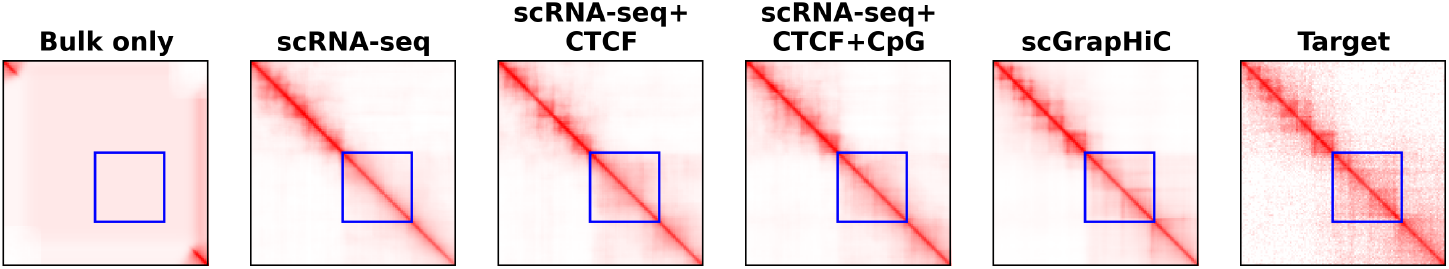
Comparison of the predicted pseudo-bulk scHi-C contact map for region chr7:32Mbp-38.4Mbp of Epiblast and PS cell type to highlight that sequence encoder methods fail to recover finer architectural features that scGrapHiC can recover, as highlighted with a blue rectangle.

We quantify these improvements using the three metrics: GD (GenomeDISCO), SCC (Stratum-Adjusted Correlation Coefficient), and TAD sim (Topologically Associating Domains Similarity). In Fig. 3, we show a violin plot of the distribution of scores (y-axis), with a mean of scores written on the blue line, across both test set chromosomes of all cell-types mentioned in the Table 1 (except the ones from Embryo stage EX15 and Brain that have been separated out for generalizability testing). Moreover, we highlight a significant change in scores with a *\** when the p-value on a Student’s t-test is ≤ 0.0001. Our quantitative evaluations show that adding structural support through CTCF score significantly improves performance, with a 4.29% improvement in GD, 52.7% in SCC, and 14.7% increase in TAD sim scores over using scRNA-seq alone. However, we do not observe a significant change in scores by adding CpG scores. We believe this happens because, at most genomic loci, the CTCF motifs overlap with CpG islands [HL13] and hence potentially contain the same information. Therefore, CpG scores consequently do not provide additional support in predicting the structure. scGrapHiC further improves performance over the sequence-based encoder baselines by 5.4% in GD, 46.6% in SCC, and 22.3% in TAD sim, demonstrating the importance of the graph deconvolution step. Furthermore, we observe that the distribution of scores for scGrapHiC is less spread out compared to other methods, highlighting its capacity to generalize to other cell types more effectively than baseline implementations.

**Abb. 3:**
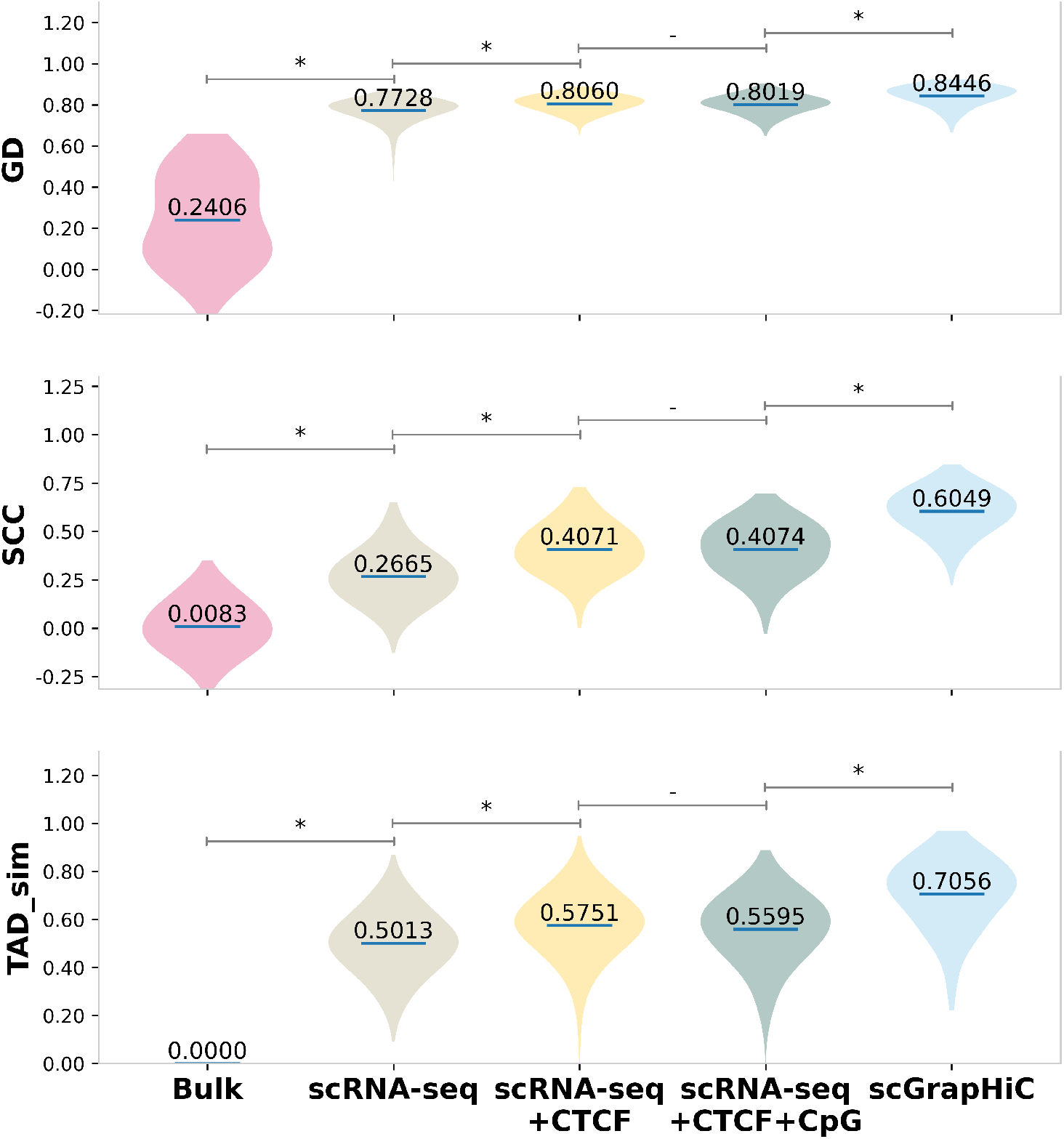
Across all three evaluation metrics, GD, SCC, and TAD sim (on the y-axis) scGrapHiC outperforms sequence encoder baselines by a significant margin, demonstrating the utility of deconvolving relevant structure from bulk Hi-C contact map.

We show qualitative and quantitative results across individual cells in the supplementary Figs. S7 and S6 highlighting scGrapHiC ‘s capacity to predict cell-specific interactions accurately. To predict cell-specific scHi-C contact maps, we find that it is crucial to embed additional support through cell-agnostic features such as CTCF and CpG scores given the weak correlation of scRNA-seq with structure. This support needs to be coupled with deconvolvution of relevant structure from the bulk Hi-C contact map through the Graph Encoder module.

### 4.2 Embryonic Stem Cell (ESC) bulk Hi-C serves as an ideal candidate for deconvolution

Given the importance of bulk Hi-C and graph deconvolution for predicting pseudo-bulk scHi-C, we investigate and identify a bulk Hi-C contact map that can serve as a generic input for an arbitrary cell-type or tissue sample scHi-C prediction. We select five candidate bulk Hi-C datasets originating from the least structurally differentiated mouse Zygote cell belonging to embryo stage E0.5 to E4.5 when mouse ESCs develop in the epiblast of the late blastocyst. These candidates were picked based on prior knowledge. It is known that the core chromatin structure is conserved throughout various cell-types and tissues. The chromatin goes through extensive lineage-specific chromatin reorganization as tissue develops from ESCs [Di15; Ou20]. A recent study on mouse embryos revealed that during embryonic development, chromatin structure shifts from a “relaxedßtate in Zygote to progressive maturation of higher-order chromatin structure in later embryo stages [Du17]. Therefore, through this experiment, we test how scGrapHiC performs when provided with the five selected bulk Hi-C contact maps.

In Fig. 4, we show a generated pseudo-bulk scHi-C contact map for the same region chr7:32Mbp-38.4Mbp of the same cell type Epiblast and PS. For all input bulk Hi-C maps, scGrapHiC can infer the higher-order genomic structure accurately; however, in the region highlighted in the blue rectangle, we find that Zygote and two-cell stage bulk Hi-C inputs struggle to recover finer genomic features, such as TADs, accurately. Beyond those stages, scGrapHiC with any bulk Hi-C input (eight cells, IMC, or mESC) produces similar outputs, with mESC producing the most accurate scHi-C outputs. Fig. 5 shows quantitative differences in performance using the five different bulk Hi-C inputs arranged in the order of their embryo stage (on the x-axis) across similarity metrics GD, SCC, and TAD sim (scores on the y-axis) in violin plot. We emphasize significant score changes compared to the previous embryo stage highlighted with a *\** (Student’s t-test p-value is ≤ 0.0001). We show the average performance of the scRNA-seq+CTCF+CpG baseline model as a gray dotted line to accentuate that even when using Zygote, the least structurally differentiated bulk Hi-C, we can achieve better scores across all three metrics. Our results show significant improvement of 1.3% in GD, 16.58% in SCC, and 11.8% in TAD sim scores. We observe an improvement of 1.5% in GD, 6.2% in SCC, and 5.1% in TAD sim scores when scGrapHiC uses ICM Hi-C contact map over the eight cells stage bulk Hi-C data. The performance with ICM and mESC is similar across GD and TAD sim, which correlates to the finding of the original study [Du17], which shows a high degree of overlap in TADs and genomic structures between ICM and mESC. Based on these results and a wider availability of high read depth ESC Hi-C data across different species, including humans and flies, we use mESC as our standard bulk Hi-C input to accurately predict cell-type specific scHi-C contact maps.

**Abb. 4:**
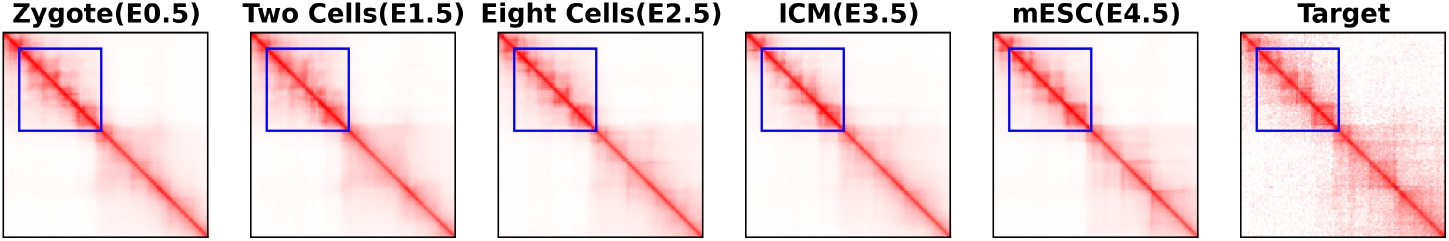
Comparison of the predicted pseudo-bulk scHi-C contact map using bulk samples from earlier embryo stages (E0.5 - E4.5) for region chr7:32Mbp-38.4Mbp of Epiblast and PS cell type to highlight that later-stage embryo bulk samples improve the quality of recovered scHi-C contact map, while the difference between ICM and mESC and eight cells is minor.

**Abb. 5:**
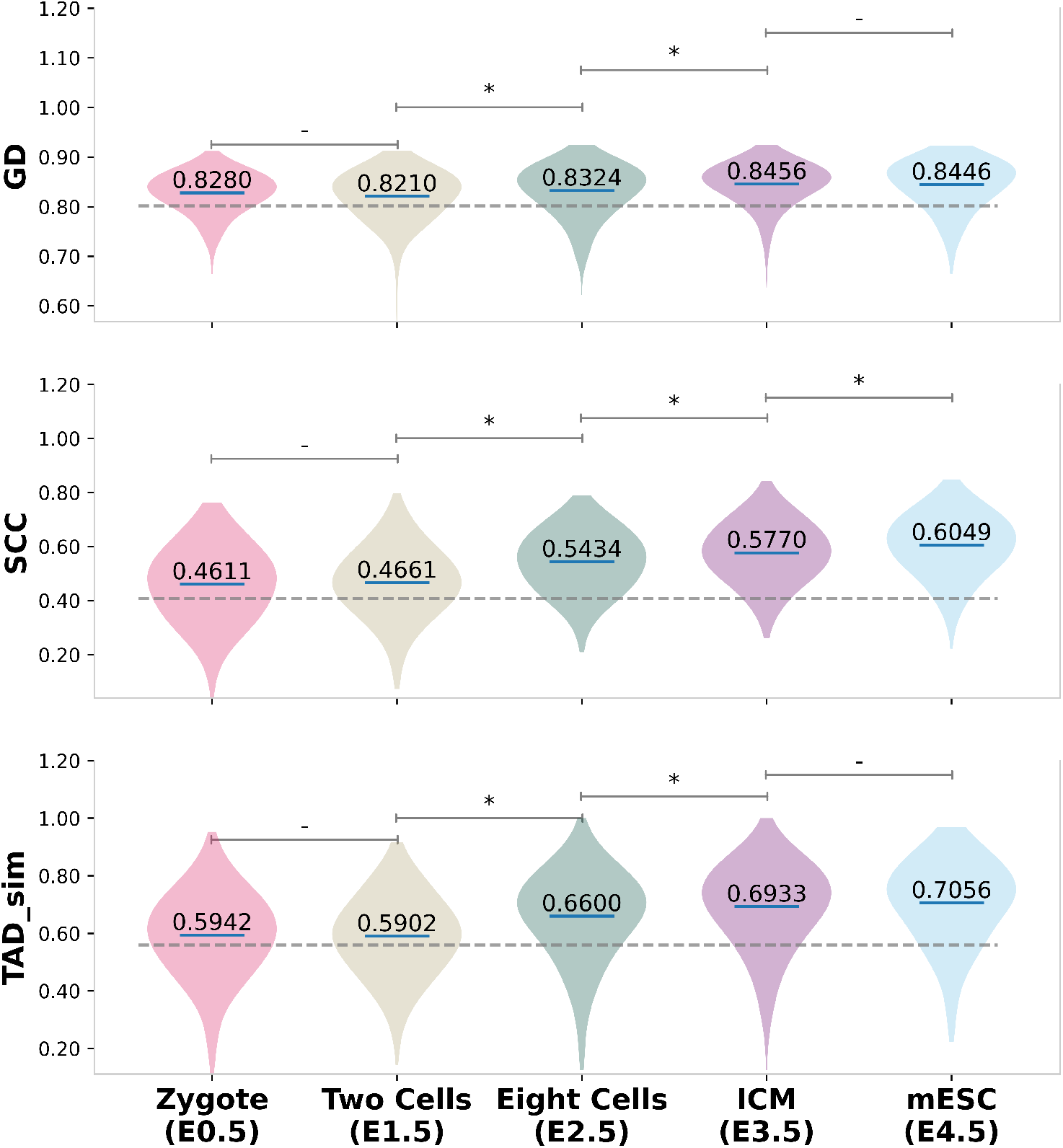
Quantitative comparison across GD, SCC, and TAD sim metrics show that using a later timepoint embryo bulk Hi-C sample improves the accuracy of the predicted pseudo-bulk scHi-C contact maps. We find that using mESC bulk Hi-C data for the deconvolution step provides the best performance. The gray dotted highlights that even with the Zygote which is the least differentiated sample, we are able to outperform the *scRNA-seq+CTCF+CpG* baseline.

### 4.3 scGrapHiC generalizes to unseen embryo stages and brain tissue samples

Next, we design an experiment to test the generalizability of scGrapHiC trained with embryo datasets E7.0 to EX05 on unseen EX15 embryo stages. We also test performance on Ex1 (Exon 1) cells from brain cortex tissue to investigate whether the model generalizes to inputs from different tissue samples. We acquired these samples from the same HiRES dataset [Li23] and separated out the cells from these cell-types or tissues from our training pipeline. As done before, the model takes cell-type-specific pseduo-bulk scRNA-seq data and cell-type-agnostic CTCF and CpG scores as feature inputs. We show the performance using mESC bulk Hi-C as input for graph deconvolution. Since chromatin goes through extensive lineage-specific reorganizations, we additionally test whether scGrapHiC can use a brain Hi-C dataset to improve performance on Ex1 and to enhance scGrapHiC ‘s capability to adapt to highly differentiated cell-types and tissue samples.

Fig. 6 compares the outputs generated by scGrapHiC for the region chr11:22.4-28.8 Mbp, which shows differential TAD structure between the embryo samples and Ex1 cell-type. As highlighted with the blue rectangle, scGrapHiC can correctly recover the TAD structure in Mix Later Mesenchyme and Early Neuron cells. While scGrapHiC mispredicts the presence of the sub-TAD structure with mESC bulk, scGrapHiC can accurately predict cell-type specific sub-TAD when provided with brain cortex bulk Hi-C. We quantify the performance of scGrapHiC on these tissue samples in Fig. 7, which, similar to previous visualizations, show a violin plot of scores across GD, SCC, and TAD sim. Fig. 7 show performance on three cell types: Mix Late Mesenchyme and Early Neurons are from embryo stage EX15 as well as Ex1 that is from the pre-frontal cortex. We observe GD and TAD sim scores similar to the average performance of scGrapHiC, shown as a gray dotted line across, for the Mix Late Mesenchyme and Early Neuron cells. However, we observe an improvement of 8% in SCC scores in Mix Late Mesenchyme cells, which can be attributed to a higher coverage with 403 cells belonging to this pseudo bulk. We observe a similar trend in the per-cell performance shown in the supplementary Fig. S5 where we also observe higher scores on average for cells with deeper coverage. In Ex1 cells, when we provide an mESC bulk Hi-C as our prior, we observe a decrease in performance compared to the average scGrapHiC scores. However, when we replace the mESC with a brain cortex prior, we find an impressive improvement of 3.2% in GD, 11.3 in SCC, and 10.9% in TAD sim scores without retraining or fine-tuning of scGrapHiC . We achieve performance scores for Ex1 cells with only 203 cells on par (or better) than cells with twice the coverage. These findings highlight that the deconvolution process capitalizes on the structure in the provided bulk Hi-C dataset to predict accurate scHi-C contact maps. The generalizability of our model suggests that we can utilize scGrapHiC to predict accurate pseudo-bulk scHi-C contact maps as long as scRNA-seq is available for any cell type or tissue samples.

**Abb. 6:**
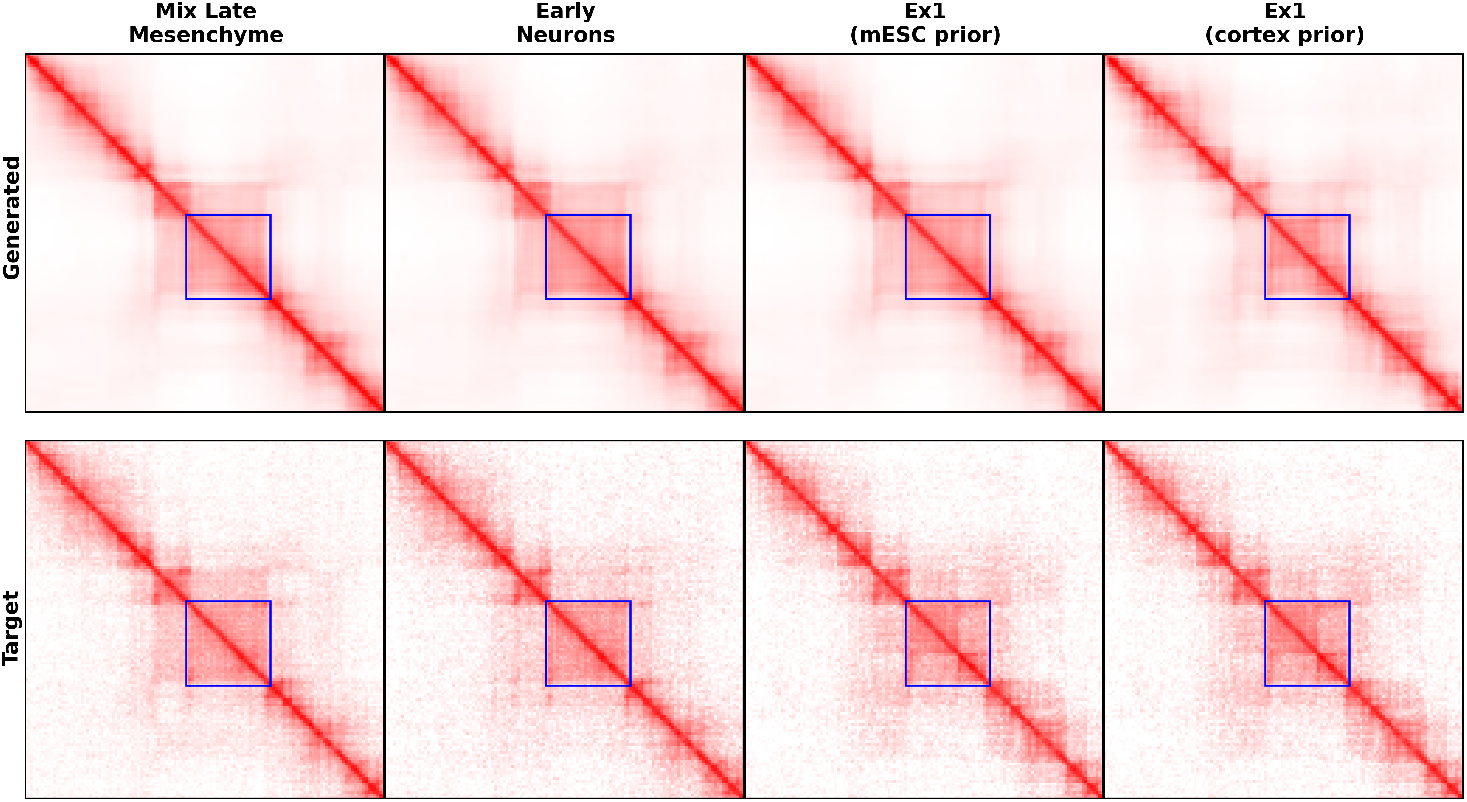
Comparison of the predicted pseudo-bulk scHi-C contact maps for held-out EX15 embryo stage cell types and Ex1 cells from brain tissue. We find that scGrapHiC can infer cell-specific TAD structures accurately and demonstrate its generalizability. For Ex1 cells, scGrapHiC can accurately infer cell-specific sub-TAD structure when provided with a brain cortex bulk Hi-C as input.

**Abb. 7:**
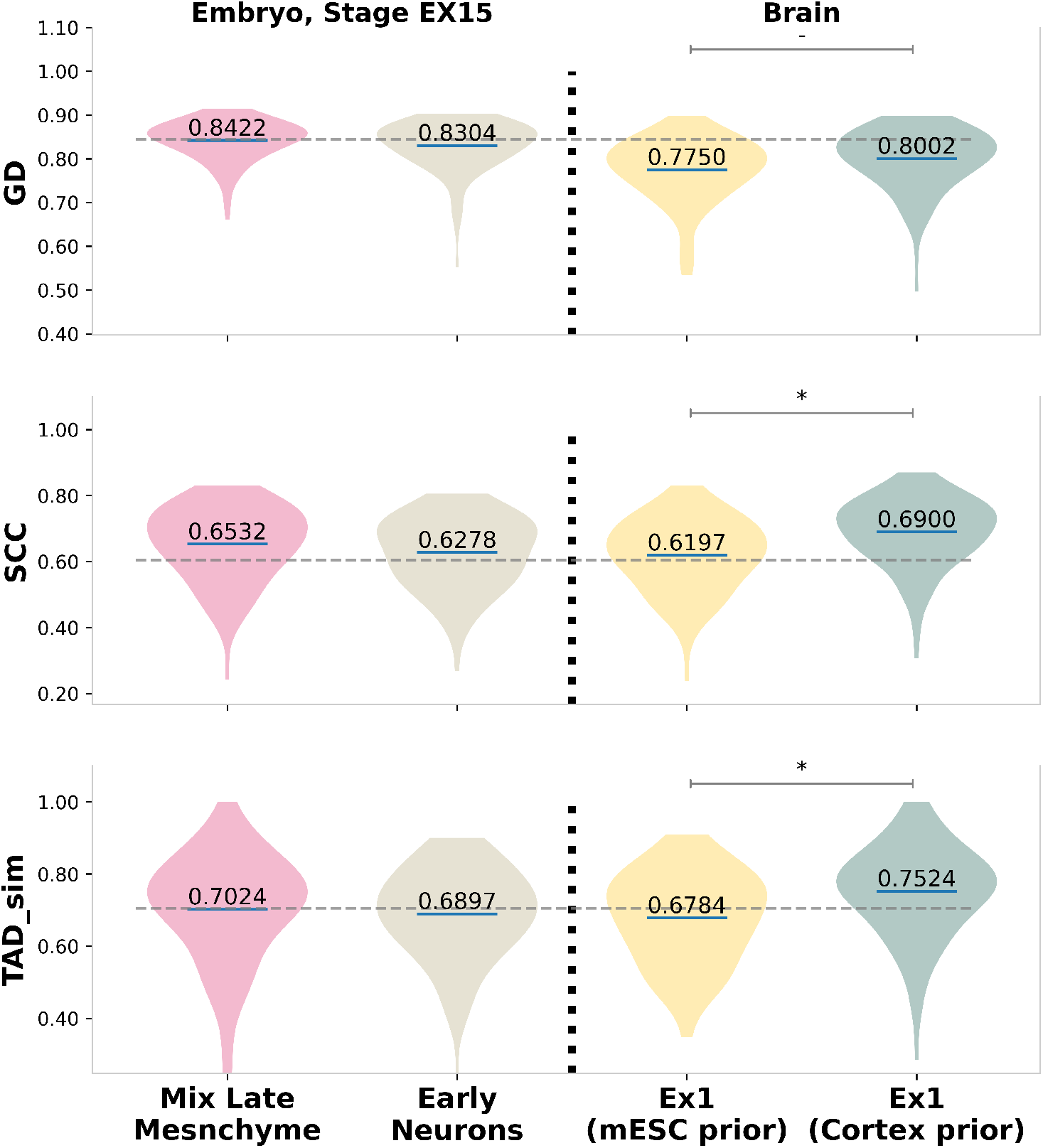
Performance of scGrapHiC on scRNA-seq data from held-out EX15 embryo stage cell types and Ex1 cells from brain tissue. Scores on the three metrics suggest that scGrapHiC can generalize well and achieve scores similar to the average scGrapHiC scores (horizontal dahsed line). Moreover, for Ex1 cells, we show that using a brain cortex bulk Hi-C contact map as input for deconvolution can significantly improve the performance, highlighting that scGrapHiC can generalize to other tissue samples when provided with the appropriate bulk Hi-C.

## 5 Discussion and Conclusion

We present scGrapHiC, a deep learning framework to predict pseudo-bulk scHi-C from scRNA-seq. It performs graph deconvolution to extracts cell-type specific scHi-C from bulk Hi-C datasets. Despite being a widely available signal, scRNA-seq is a challenging dataset to map to scHi-C because the genomic structure does not directly correlate to gene expression profiles. We can simplify this mapping task by relying on sequential, cell-agnostic structural priors provided through CTCF motifs and CpG scores. However, introducing bulk Hi-C as an additional prior on structure allows us to deconvolve cell-specific scHi-C using scRNA-seq as a guide. This bulk Hi-C prior allows us a fine control on what structure scGrapHiC deconvolves from, and we have shown that when predicting for Ex1 cell type, providing a cerebral cortex bulk Hi-C boosts the performance substantially over using mESC as a prior.

Given the technical challenges and high sequencing costs of scHi-C, our method scGrapHiC presents a robust alternative to augment scRNA-seq with structural information to help researchers disentangle the complex relationships between gene expression and the genome structure. scGrapHiC has more potential than existing Hi-C prediction methods since we rely on scRNA-seq that is a more established and frequently used experimental technique than scATAC-seq. In the future as the coverage of both scRNA-seq and scHi-C evolves, we can reduce the number of cells required per pseudo bulk (less than 190) sample and hence would allow us to study the structure of rarer cell types. Lastly, as more scRNA-seq and scHi-C co-expression data becomes publicly available, we will evaluate our model on data from different species [Zh23] and more tissue samples.

---

**Abb. S1:**
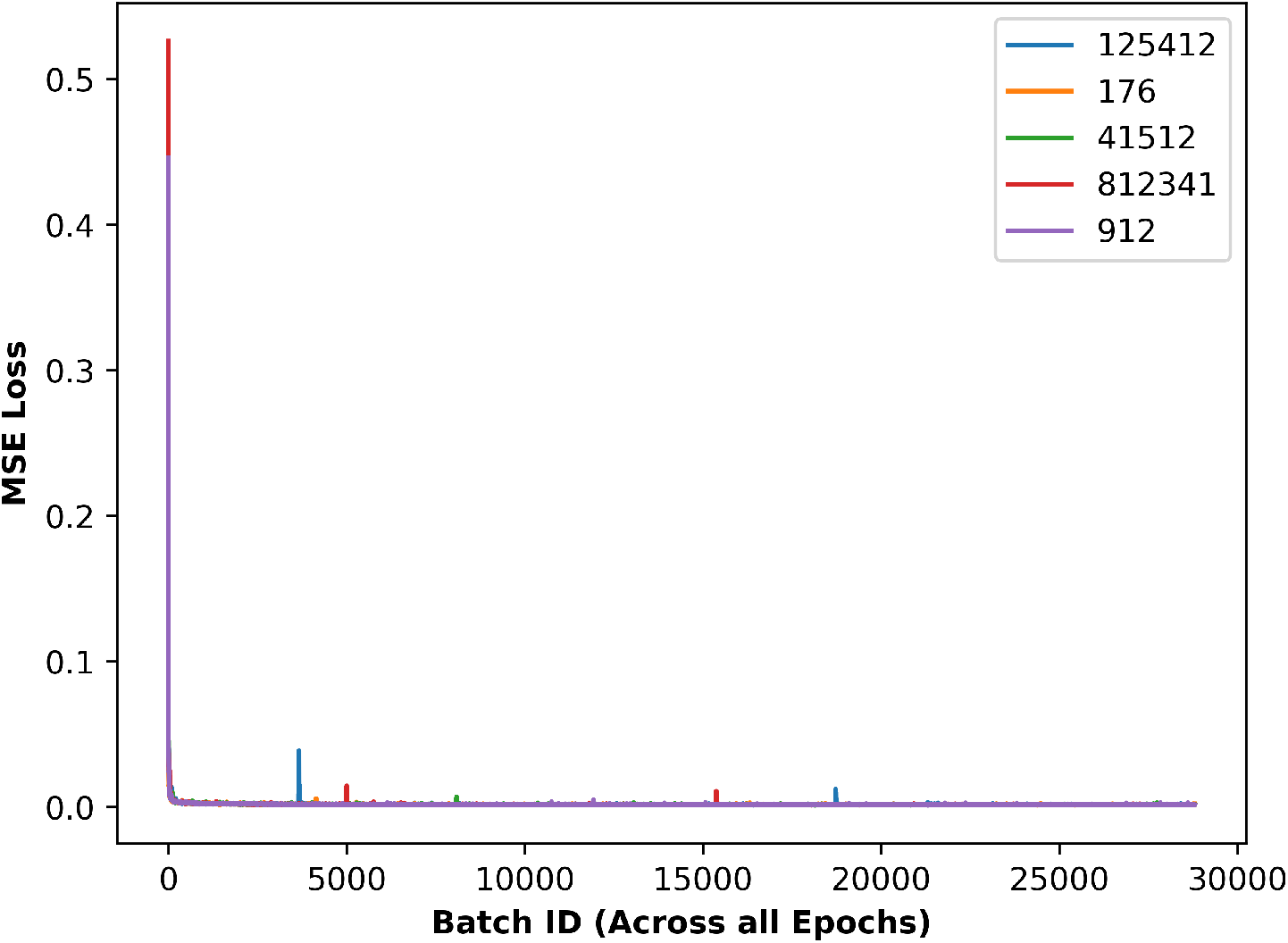
We show the loss curve of scGrapHiC of 5 different random seeds and find that the model converges to a very similar loss highlighting scGrapHiC ‘s stability to randomized weight initialization. For rest of the evaluations, we choose another random seed of 40 and show rest of the results on that seed.

**Tab. S1:** This table mentions all the hyperparameters, what their purpose is and refers to the appropriate figure/table explaining the tuning process.

|  | Value | Explanation | Reference |
| --- | --- | --- | --- |
| Library Size | 25000 | The library size we aim to simulate through our library size normalization pre-processing step. | Fig. S2 |
| Soft Threshold ($t$) | 1 | Soft threshold value we use for smoothing the scHi-C contact matrices | Fig. S3 |
| Positional Encodings Dimensions ($k$) | 16 | Top $k$ components that we select from the eigenvectors of the bulk Hi-C as positional encodings | Fig.S4 |
| Supporting Features | True | Whether we use the supporting structural features CTCF and CpG scores as additional node features | S4 |
| Number of cells cutoff | 190 | We filter our cell-types during pseudo-bulk that have less than 190 cells. | Fig. S5 |

**Abb. S2:**
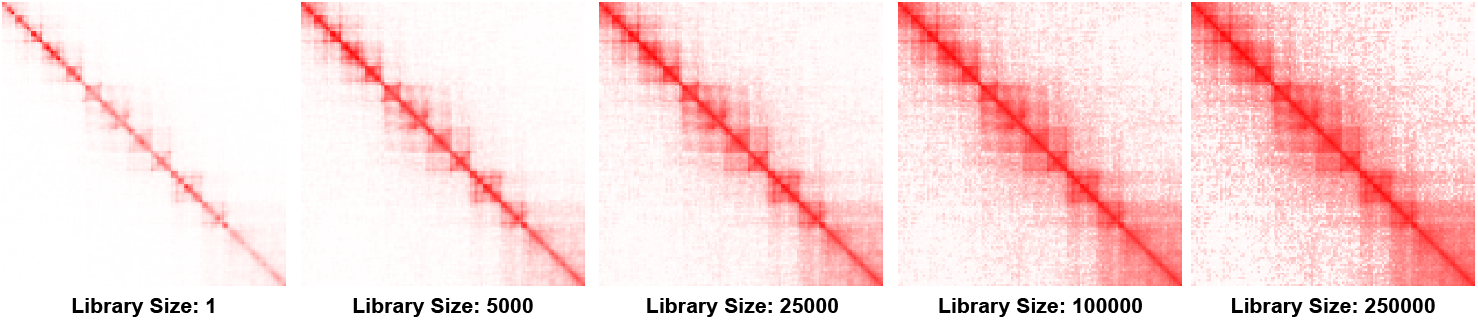
Tuning of the library size parameter, we that using a value of 25000 allowed us to enhance the structure while keeping the background noise low.

**Abb. S3:**
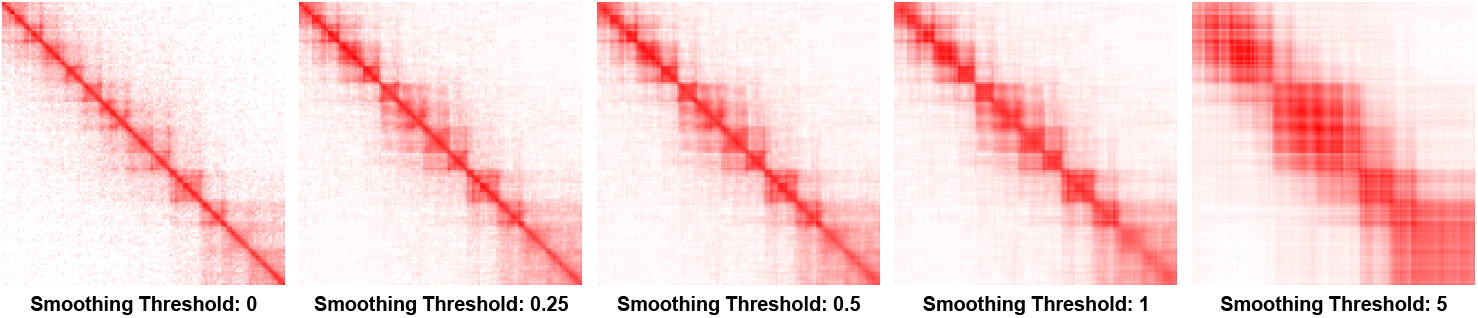
Tuning of the Hi-C smoothing soft thresholding parameter, and we found that using 0.25 ensures that we are able to supress the background noise while preserving the hierarchical organization of the genome.

**Abb. S4:**
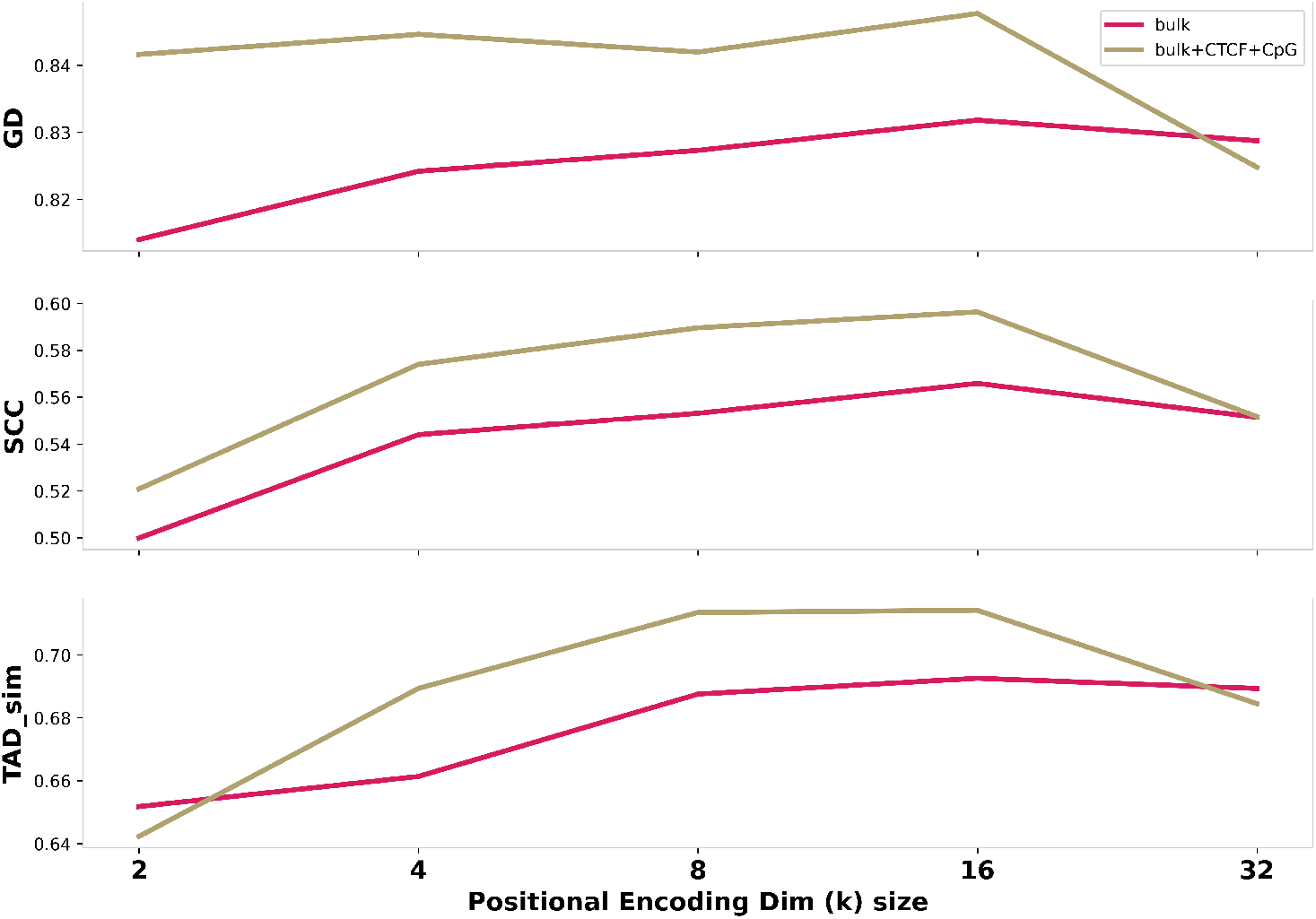
Positional encoding vector of size 16 found through hyperparameter tuning on different values of *k*

**Abb. S5:**
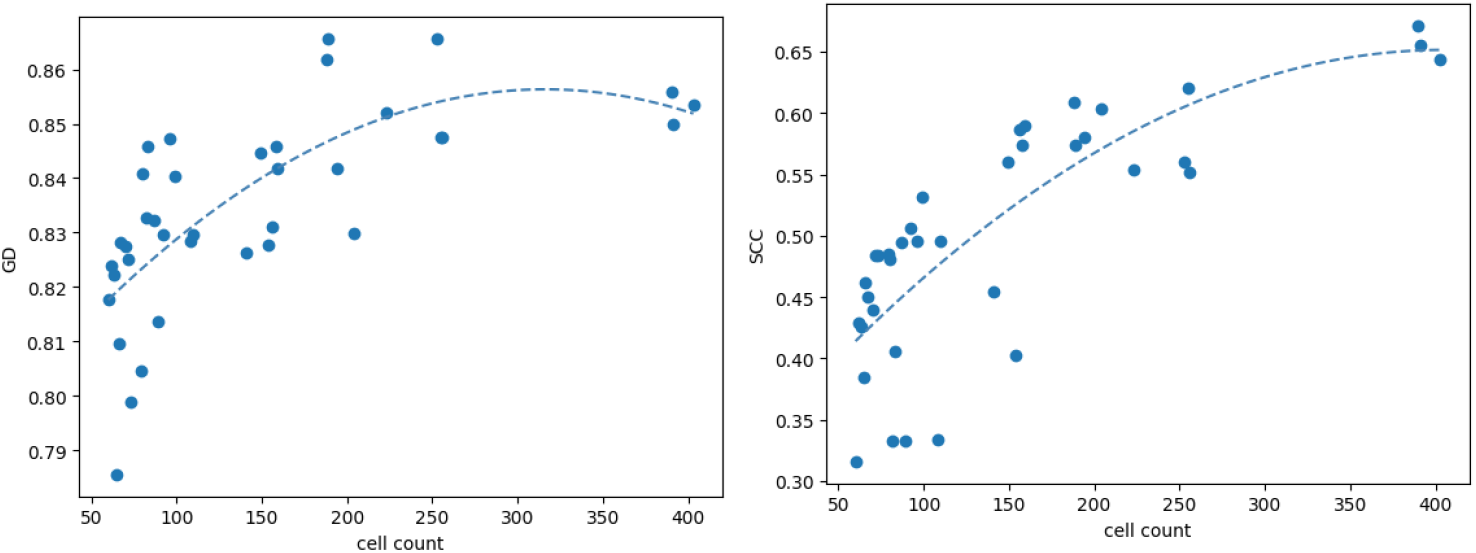
Adjusting for the pseduo-bulk number of cells cutoff parameter shows that the performance degrades non-linearly as the number of cells per pseudo-bulk go down. Suggesting a drastic loss of structure and cell-identifying features making it challenging for the current setup to realibly predict scHi-C contact maps.

**Abb. S6:**
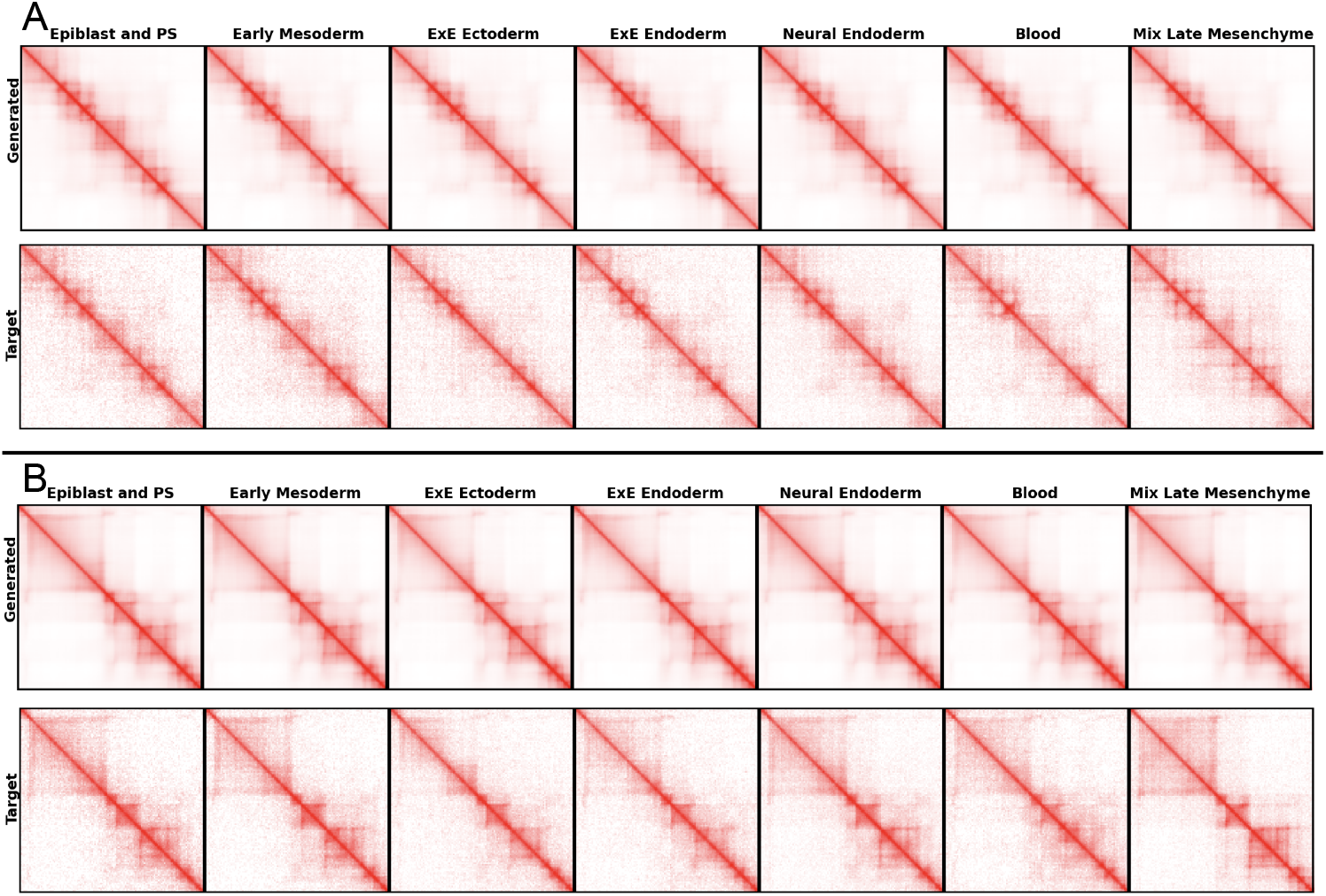
We show two different regions from Chr 7 in **A** and Chr 11 in **B**. We find that scGrapHiC is able to accurately infer complex chromatin hirearchical strucures accurately. However, in **B**, we also find that scGrapHiC mispredicts the presence of a sub-TAD structure in blood cell-type which we believe can be attributed to sparse coverage either in the scRNA-seq or scHi-C or both for this cell-type.

**Abb. S7:**
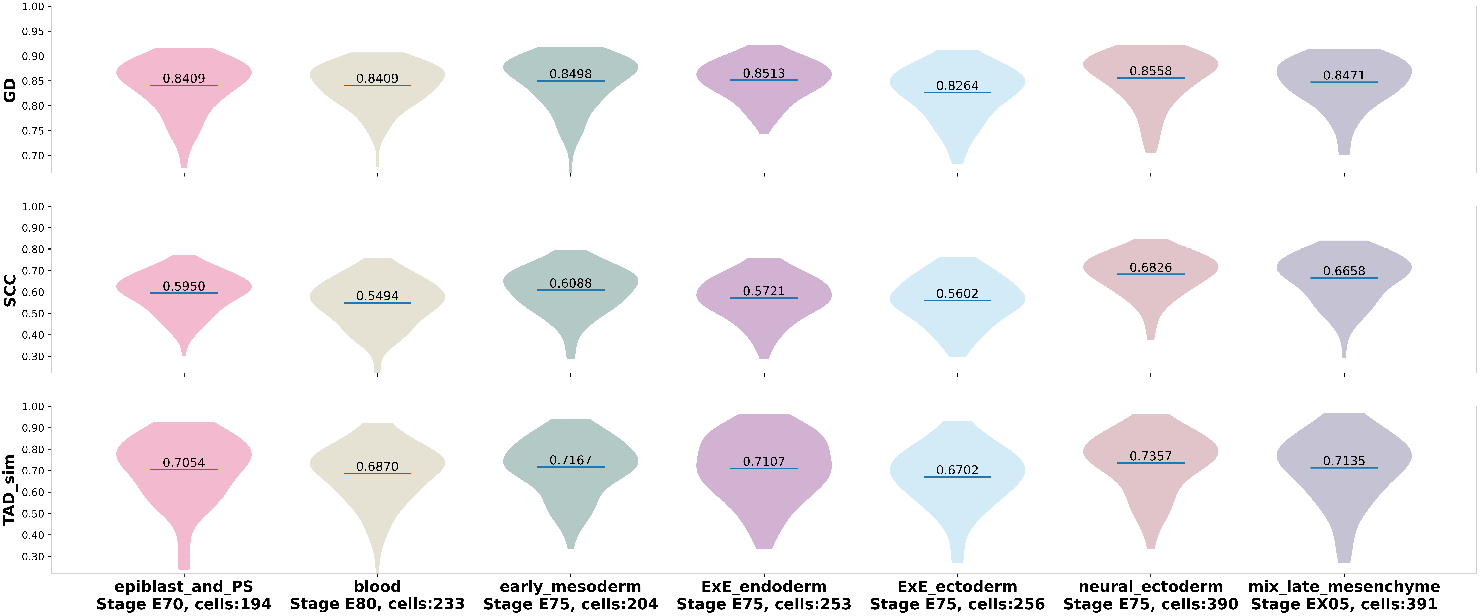
We show the scores on each cell-type individually across GD, SCC and TAD Sim metrics. We find that the scores stay fairly consistent however, the cells with lower coverage tends to get less scores in all three metric. This correlation highlights the inherent limitation of working with sparse single cell datasets.

## Footnotes

3 Reliance on the metadata file is optional since we can get the same pseudo bulks through other cell clustering approaches such as Metacells [Ba19]

## Notes

### Competing Interest Statement

The authors have declared no competing interest.

